# Building a tRNA thermometer to access the world’s biochemical diversity

**DOI:** 10.1101/2020.07.01.179846

**Authors:** Emre Cimen, Sarah E. Jensen, Edward S. Buckler

## Abstract

Because ambient temperature affects biochemical reactions, organisms living in extreme temperature conditions adapt protein composition and structure to maintain biochemical functions. While it is not feasible to experimentally determine optimal growth temperature (OGT) for every known microbial species, organisms adapted to different temperatures have measurable differences in DNA, RNA, and protein composition that allow OGT prediction from genome sequence alone. In this study, we built a model using tRNA sequence to predict OGT. We used tRNA sequences from 100 archaea and 683 bacteria species as input to train two Convolutional Neural Network models. The first pairs individual tRNA sequences from different species to predict which comes from a more thermophilic organism, with accuracy ranging from 0.538 to 0.992. The second uses the complete set of tRNAs in a species to predict optimal growth temperature, achieving a maximum *r*^2^ of 0.86; comparable with other prediction accuracies in the literature despite a significant reduction in the quantity of input data. This model improves on previous OGT prediction models by providing a model with minimum input data requirements, removing laborious feature extraction and data preprocessing steps, and widening the scope of valid downstream analyses.

## INTRODUCTION

Environmental temperature affects every biochemical reaction within an organism, from spontaneous protein folding to complex metabolite catalysis. Tools that infer an organism’s optimal growth temperature from genomic sequence have potential biological and economic implications and can improve understanding of how both individual proteins and whole organisms adapt to their environment. However, experimentally identifying the true optimal temperature of every newly-discovered microorganism is unfeasible due to the sheer number of prokaryotes that have been identified and the difficulty in isolating and culturing many prokaryotic species. Predicting an organism’s optimal growth temperature based on physical characteristics of the genome is one way to determine optimal temperature without needing to successfully culture a new species.

Temperature has a significant effect on cell biochemistry. In general, cellular processes speed up as temperature increases, but extremely high temperatures can also denature proteins and negatively affect biochemical reactions. Proteins function best within a specific temperature window that maximizes enzymatic reaction rate without denaturing the protein, and most enzymes have evolved within an optimal temperature range that is closely tied to environmental temperature (1). Few enzymes show optimal activity more than 10°C above or below the optimal growth temperature of the host organism (1, 2). Maintaining catalytic function at extreme temperatures requires specific changes in genome composition, and many studies have identified differences between thermophilic and mesophilic genomes (1, 3–11). Thermophilic proteins tend to have more hydrophobic residues, disulfide bonds, and ionic interactions to pack amino acid residues closely together and prevent protein unfolding, while psychrophilic (cold-adapted) proteins require fewer strong interactions between amino acid residues (2). Significant shifts in genome composition have also been correlated with environmental temperature conditions, affecting genome features like GC content, codon bias, and amino acid frequency (12, 13).

Because physical changes in DNA, RNA, and protein composition have been correlated with optimal growth temperature, multiple approaches have been taken to predict OGT from a combination of these features. Aptekmann *et al.* found a positive correlation between G C content of tRNA regions and OGT, but not in the overall genome and found a positive correlation between information content and optimal growth temperatures in Archaea (12). Li et al. implemented a machine learning workflow in order to predict the OGT of microorganisms and enzyme catalytic optima from 2-mer amino acid composition and investigated a wide range of regression models (5). Sauer and Wang used multiple linear regression to predict the OGT of prokaryotes from genome size and tRNA, rRNA, open reading frame, and proteome composition. (13). Ai et al. targeted a problem related to protein thermostability and classified thermophilic and mesophilic proteins using support vector machines and decision trees (6). Similarly, Capaldi et al. predicted bacterial growth temperature range based on genome sequences with a Bayesian model (1).

These previously-published approaches to predicting organism OGT tend to use a combination of genome and sequence features that cover the entire Central Dogma of molecular biology. Such models can be useful when the end goal is to predict OGT for a newly-identified species, but they have limited downstream applications when the end goal is understanding how cellular components adapt or evolve under different temperature conditions, because using many cell features to predict OGT then statistically confounds downstream analyses involving the same cellular components. Optimal growth temperatures predicted with a model requiring amino acid composition as input, for example, cannot be used in subsequent analyses investigating how proteins evolve under different temperatures – the protein evolution results would be confounded with the initial OGT prediction. To address this, we set out to create a model that predicts prokaryote OGT using a minimal set of input data. We developed a Convolutional Neural Network (CNN) model to classify and predict prokaryote OGT using only tRNA sequences as input data. We chose to focus on tRNA sequences because tRNAs are a vital and universal part of life, and RNA base pairing chemistry is known to be affected by temperature (14). Because the model only uses only tRNA features, it is possible to use predicted optimal temperatures from this model in downstream analyses evaluating the effects of temperature on other cellular components, including protein and genomic features. Essentially, with only ~2400bp of sequence (~0.1% of the genome), this CNN model can predict OGT as well as summary data from the entire rest of the genome. Because it uses fewer features than previous OGT-prediction models and does not require feature extraction, it is also easier to use and removes researcher bias in selecting features.

## MATERIAL AND METHODS

### Data collection and distribution

A list of species for which optimal growth temperature has been determined was obtained from Sauer *et al.* and all existing genome assemblies were downloaded for each species, resulting in an initial set of 36,529 Bacteria and 276 Archaea genomes, with optimal growth temperatures ranging from 4 to 103°C (13). tRNA sequence and positions were predicted for each genome using tRNAscan-SE (version 2.0.3; 15, 16). rRNA sequence and positions were predicted for each genome using barrnap (version 0.9; 17).

A single genome was selected for each species, and only tRNA sequences from that genome were used in the CNN predictions. Genome assemblies were considered low-quality and removed if 16S, 23S, and 5S rRNA sequences could not be predicted by barrnap. For species with multiple remaining genome assemblies, the assembly with the highest number of predicted tRNA sequences was chosen as the single ‘best’ assembly. Selecting genome assemblies in this way resulted in genomes for 165 unique archaea species and 2375 unique bacteria species. However, the distribution of optimal temperatures for this species set was highly skewed, with nearly 40% of archaea and nearly 70% of bacteria genomes having predicted optimal temperatures of 28, 30, or 37°C. These three temperatures are common prokaryote culture temperatures so we decided to remove observations at these temperatures to reduce the bias and help balance the dataset, since we could not be sure that these were true adaptive growth temperatures rather than culture temperatures. After removing species with OGT listed at 28, 30, or 37°C, the dataset contained 683 bacteria species with 41,853 predicted tRNAs and 100 archaea species with 5,474 predicted tRNAs.

### Prediction Method

We used Convolutional Neural Networks (CNNs) for OGT prediction. CNNs are neural network (NN)-based machine learning models with at least one convolutional layer. They can predict both categorical (classification) and continuous (regression) targets depending on the configuration of downstream layers. When the input is a continuous or discrete signal such as images, sensor data, or base pair sequence, CNNs are useful because they automatically extract features from the input and can capture both local and global features. Moreover, thanks to weight sharing at the convolutional layers and downsampling at the pooling layers, CNNs have fewer parameters that need to be learned than regular NNs. Thus, CNNs require less training data and have lower risk of over-fitting. Additionally, other machine learning models have no prior knowledge of how input values are organized, and are not able to take advantage of the relative positions within a sequence, (i.e., they cannot determine consecutive bases in a sequence). CNN architectures naturally have this prior neighborhood knowledge.

We set two prediction problems in this study. One is a binary classification model that can take two tRNA sequences from different organisms as input. This classification model predicts which tRNA sequence in the pair comes from a microorganism with higher optimal growth temperature, and requires only a single tRNA from each genome for the classification. The second model uses a CNN regression model that predicts the optimal growth temperatures of bacteria and archaea. In both cases, a CNN model was chosen to allow automatic feature extraction. This made it possible to use tRNA sequences as direct input and did not require manual extraction of hundreds of sequence features or assessment of the correlation of each individual feature with OGT. In both models, tRNA sequences were obtained as described above, then one-hot encoded. Because not all tRNA sequences are the same length, shorter sequences were padded with zeros to produce a final 4xL matrix of 0s and 1s, where *L* is the length of the longest tRNA in the input data.

### Models

#### Temperature Classifier Model based on Individual tRNA sequences

In the first model, we built a CNN classification model that can take paired sets of tRNAs and predict which tRNA belongs to a microorganism with higher optimal growth temperature. Before presenting our classifier model, in Equation 1 and 2 we introduce the data structure. In Equation 1, dataset *D* has *N* microorganisms. Each microorganism m_i_ has a corresponding OGT t_i_. In Equation 2, because each organism in the dataset has more than one tRNA, *n_i_* is the number of tRNAs in microorganism *m_i_*. Since there are a different number of tRNAs in each microorganism, *n_i_*’s are different for each *m_i_*. 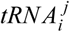 is the *j^th^* tRNA of *i^th^* microorganism *m_i_* and, it is an instance in *R*^4×*L*^ space. The input space is 4 × *L* dimensional because there are 4 base pairs (one-hot encoding), and the length of the longest tRNA in the dataset *D* is *L*.

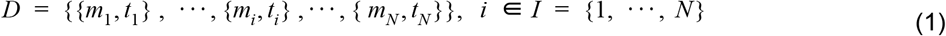

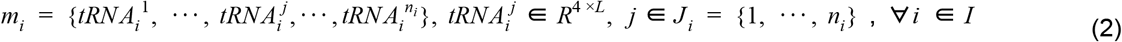

To predict which tRNA comes from an organism with higher OGT, we built a CNN classifier with two branches, each of which was fed an input tRNA sequence. Each branch starts with at least one pair of convolutional and pooling layers. After the last pooling layer, branches are flattened and merged. Convolutional and dense layers use a Rectified Linear Unit (ReLU) activation function. Two output nodes at the end are activated with the Softmax function which provides class probabilities. We predict the output as a binary label that indicates whether the first branch input or the second branch input has a higher OGT. The loss function of the model is the categorical cross entropy, and it is minimized with the Adam optimizer. Parameters of the model (e.g. layer sizes, number of layers, optimizer parameters and dropout rate etc.) were selected with hyper-parameter optimization. To train and test our classifier model, we created a dataset of tRNA pairs from the original dataset given in Equation 1 and Equation 2. In the classification dataset each sample is in the form of {{*tRNA*1, *tRNA*2}, *y*}, where *y* ε {0, 1} is the class label. We used tRNA pairs only if their microorganism OGTs were different by at least 1°C (Figure 1A).

**Figure 1.**
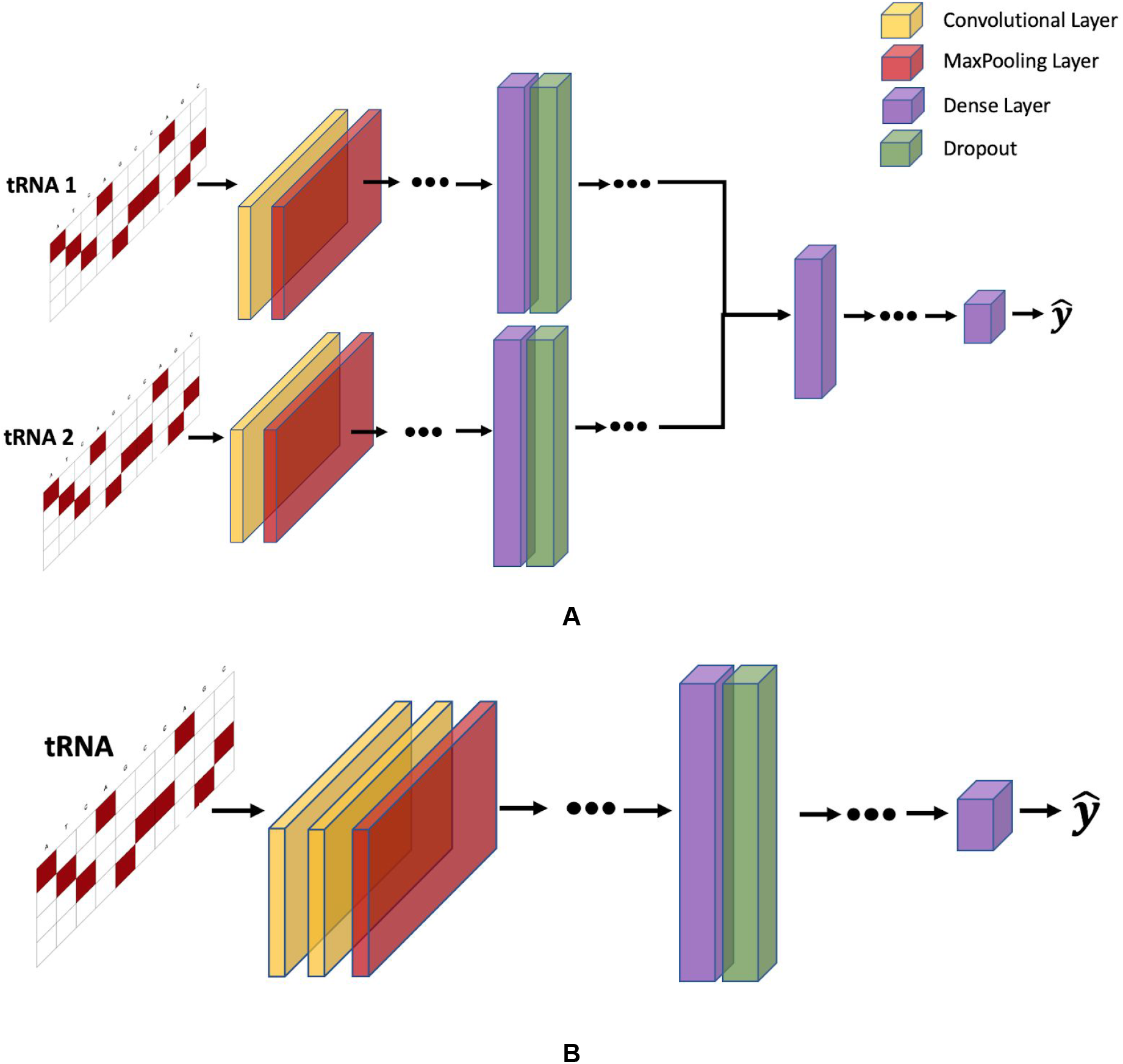
OGT prediction models. Input tRNA sequences are one-hot encoded and padded with 0s to make the matrix. A) The general structure of the temperature classifier model. The model has two channels. Both channels are fed a tRNA. Each channel starts with a convolutional layer and a maximum pooling layer. The sequence is flattened and passed to two fully connected and dropout layers. The two channels are concatenated and then passed to fully connected layers. The output layer has two neurons for binary classification. B) The general structure of the regression model. Input layer is followed by two convolutional layers and a maximum pooling layer. Then, data are processed through fully-connected dense layers, resulting in a single OGT prediction for each tRNA. This figure shows the general structure only, and the exact number of layers are selected with hyper-parameter optimization. Selected hyper-parameters are provided in the supplemental files.

#### Species OGT Predictor Model

The CNN regression model to predict species’ OGT starts with two convolutional and subsequent maximum pooling layers. After the last pooling layer there is a single flattening layer before multiple fully-connected dense layers. The activation function of the convolutional and dense layers is Rectified Linear Units (ReLU; Figure 1B). The output node is a continuous variable and is activated linearly. The loss function of the model is the mean squared error, and it is minimized with Adam optimizer. In Figure 1, we provide the general structure of the models. Dots mean the model may have more layers of the given type. The number of layers for each model is selected with the hyper-parameter optimization.

To train and test the regression model, we created a dataset as in Equation 1 and Equation 2, where each sample is in the form {*tRNA, t*}. *t* is OGT of the related tRNA. We trained the CNN model by considering each tRNA as independent of all other tRNAs in the organism. Once we trained the model, we had *n_i_* tRNA-based OGT predictions for the microorganism *m_i_*; one for each tRNA. We determined the OGT prediction of the species as a whole by calculating the median of all tRNA-based OGT predictions. In Equation 3, 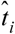 is the OGT prediction of *i^th^* microorganism *m_i_* and, 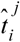 is the OGT prediction of *j^th^* tRNA of *i^th^* microorganism.

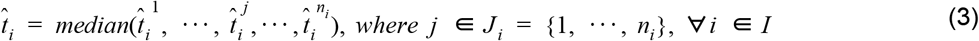

### Hyper-parameter Optimization

Selecting the optimal combination of hyperparameters is important because hyperparameter values have a significant effect on the performance of CNN models. There are a large number of hyper-parameters and the possible values of each results in millions of potential hyper-parameter combinations. In this study, we used Bayesian optimization to determine parameter values for layer size, the number of layers, the number of filters, kernel size, pooling size, strides, dropout rate, batch size, and beta1, beta2, learning rate of the Adam optimizer. The search space for each hyper-parameter is listed in Supplemental Table 1.

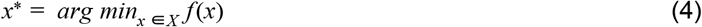

Selecting the best set of hyper-parameters *x*\* from the possible selection set *X* is defined in Equation 4, where *f*(*x*) is the loss (e.g. mean squared error, mean absolute error, classification error) on the validation set. The optimization-based approach that we use maximizes the expectation of the improvement (EI) on the performance. There are several ways to estimate EI, and we used a tree-structured Parzen Estimator (TPE) in the Hyper-opt python package to find *x*\*, which provides the highest EI (18, 19). We ran the algorithm for 50 iterations to select the best combination of hyperparameters to use in each model.

### Data splitting procedure

To evaluate the performance of each proposed model, we investigated two scenarios. First, we split the species randomly into training, validation and test sets. Second, we controlled for evolutionary relatedness and split data according to phylogenetic distance.

#### Random Split

Model evaluation commonly uses *k*-fold cross-validation or a random split of the training, test, and validation data sets. In the first part of the computational results, we held out 5% of all species as a validation set to fine-tune hyper-parameters and then used the rest of the species for 5-fold cross-validation: 76% for training, and 19% for testing in each iteration of the model. Hyper-parameter optimization was done once during the first iteration, and hyperparameters were selected according to performance on the validation set. We repeated the whole set of tests five times.

#### Phylogenetic distance split

Random train-test data splits work well for prediction models, and previous OGT-prediction models use a random training/test split to evaluate their model (5, 6, 9, 13). However, the random training/test split does not account for evolutionary relatedness between species. Previous machine learning studies have found that similarity between individuals can provide overly-optimistic results because the training and test data sets may contain closely-related species. Washburn *et al.* (20) considered this situation in the prediction of mRNA expression levels, and found that machine learning models trained without taking evolutionary history into account were able to recognize species similarity and use it to inform predictions. Additionally, Washburn *et al.* states that ignoring shared evolutionary history can exaggerate model performance and possibly lead researchers to conclude that certain model features are important, when in fact, the model test set is contaminated by similar observations that are present in both the training and test sets. In fact, if the aim of using a predictive model is only to predict accurate OGTs/class labels, splitting training and test sets randomly is valid. However, if researchers would like to obtain biological insights from the model (i.e. to answer questions like which tRNA properties are correlated with high/low OGT), a dataset split by phylogenetic distance will provide better insights. A model trained using a phylogenetic distance train/test split is also likely to be more transferable to other problems. This is because by constraining the model and removing all information from phylogenetically related species, we push the model to extract other rules from the sequence that are more transferrable.

To account for evolutionary relationships across species, we tested each model with a phylogenetically-informed training/test split. A simple phylogenetic relationship between species was calculated as the relative relationship between species based on species’ taxa id and the NCBI Common Tree (21, 22). The tree was converted to a simple distance matrix using the ‘ape’ R package (23). The phylogenetic distance matrix was used to split species into clusters. For this purpose, we applied hierarchical clustering to species by minimizing the Ward variance (24). After preserving 5% of the species for the validation set, the rest of the species were split into 10 clusters, 8 of which were used as a training set and 2 of which were used as the testing set. This procedure was repeated 5 times, and each cluster was included in the test set only once. We repeated the whole set of tests five times.

#### Model attention

To determine how the importance assigned to these stem structures affects model predictions, we selectively mutated each set of paired bases in the tRNA structure for 55 E. coli tRNAs for which structure information is available (26). For each paired nucleotide we mutated the original base to all three other nucleotides (e.g. G → A, T, and C in turn) to disrupt a single Watson/Crick base pairing interaction within the tRNA. This resulted in a set of 5,793 new tRNA sequences, each with a single nucleotide change that disrupted one set of pairing interactions. We then predicted OGT for these new sequences and compared predictions to OGT predictions for the original E. coli tRNA sequences.

#### Software and Computation Power

We implemented the proposed prediction method in Python 3.6 using Keras (2.2.5) to build and train the CNN model. For the distance split, species were clustered hierarchically with Sklearn (0.21.2) to find clusters related to phylogenetic distance. Tests were carried out on a computer with a GeForce RTX 2080 GPU, 64 GB RAM, and Intel(R) Core(TM) i7-7800X CPU running at 3.5 GHz.

## RESULTS

### Model 1: Temperature classifier based on individual tRNA sequences

We first built a model that uses a single tRNA to distinguish which of two species comes from a higher optimal growth temperature environment. Data was split into training and test sets either randomly or with phylogenetically-informed distance splitting as described above. All tests were repeated five times and results represent the average and the standard deviation of these runs.

For both the random and phylogenetic data splits we calculated accuracy results when temperature differences are greater than 0 C, 5 C, 10 C, 20 C and 30 C. For bacteria genomes, including phylogenetic information as well as the tRNA sequence improved predictions (Figure 2A). Interestingly, the model results showed that in archaea genomes, a single tRNA pair contains enough signal to distinguish between genomes from species adapted to different optimal growth temperatures, even when phylogenetic relationships between species have been deliberately removed (Figure 2B). Unsurprisingly, the model performs better when there are larger temperature differences between the species being compared. It is possible that the model learned species OGT, rather than sequence features related to OGT, so we also verified that the model was not just predicting the same direction for a given species in the results (e.g. was not always predicting “lower OGT” for a given species).

**Figure 2.**
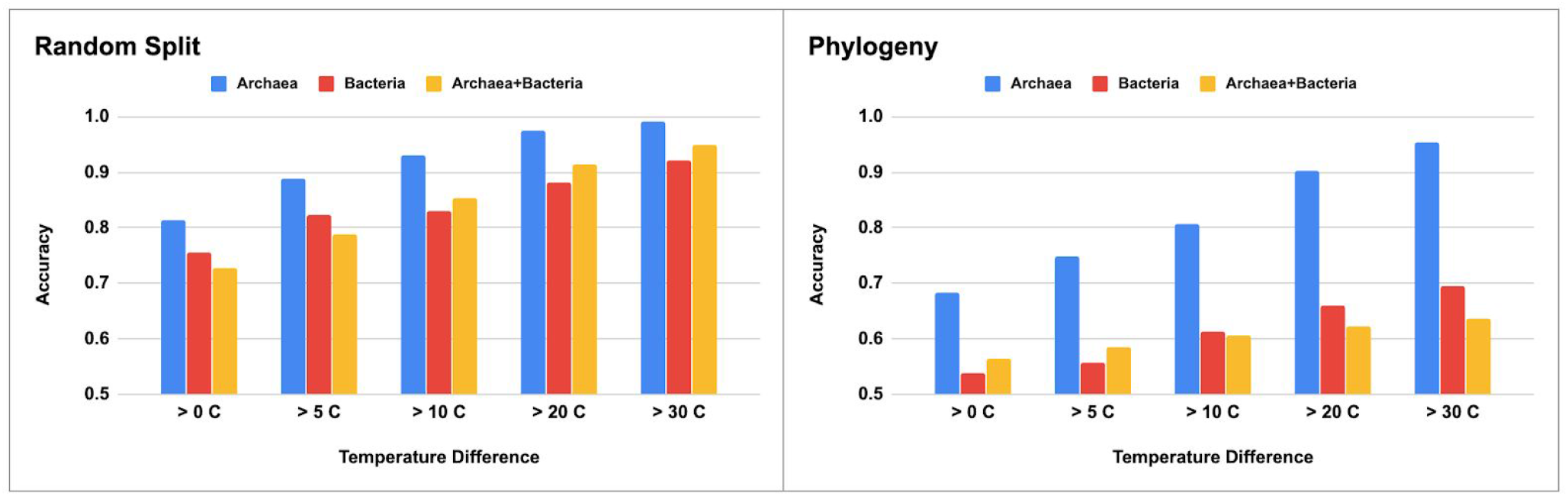
CNN classification results for models built for each phylogenetic domain and with either A) randomly split training and test datasets or B) phylogenetically-informed training and test datasets.

### Model 2: Species OGT prediction

We next asked whether it was possible to use information from all tRNAs within a species to predict overall species OGT using a regression CNN model. As with the classification model, prediction accuracy was compared using both random and phylogenetically-informed data splits. Data was split into hyperparameter validation, training, and test sets as described in the classification model. Root mean squared error (RMSE) and the coefficient of determination (*r*^2^) were used to evaluate model performance for both the random and phylogenetic distance split models. A good model will have both low RMSE and high *r*^2^. Low RMSE indicates how closely the model can pinpoint a species’ OGT, while high *r*^2^ indicates that most of the variance in the true OGT dataset is explained by the OGT predictors. RMSE and *r*^2^ were compared for each domain individually as well as the combined domains (Table 3, Figure 2).

**Table 1.** Classification model accuracy performance with Archaea, Bacteria, and combined dataset.

| Temperature Difference | Archaea |  | Bacteria |  | Archaea + Bacteria |  |
| --- | --- | --- | --- | --- | --- | --- |
|  | Random | Phylogenetic Distance | Random | Phylogenetic Distance | Random | Phylogenetic Distance |
| > 0 °C | 0.814±0.011 | 0.684±0.025 | 0.756±0.001 | 0.538±0.001 | 0.728±0.023 | 0.564±0.029 |
| > 5 °C | 0.889±0.010 | 0.749±0.029 | 0.824±0.002 | 0.557±0.012 | 0.788±0.030 | 0.585±0.039 |
| > 10 °C | 0.930±0.005 | 0.807±0.025 | 0.830±0.005 | 0.612±0.016 | 0.853±0.033 | 0.605±0.0512 |
| > 20 °C | 0.974±0.004 | 0.902±0.011 | 0.882±0.005 | 0.660±0.024 | 0.914±0.032 | 0.622±0.063 |
| > 30 °C | 0.992±0.002 | 0.954±0.008 | 0.921±0.007 | 0.694±0.035 | 0.950±0.037 | 0.636±0.071 |

**Table 2:** Classification model F1 score with Archaea, Bacteria, and combined dataset.

| Temperature Difference | Archaea |  | Bacteria |  | Archaea + Bacteria |  |
| --- | --- | --- | --- | --- | --- | --- |
|  | Random | Phylogenetic Distance | Random | Phylogenetic Distance | Random | Phylogenetic Distance |
| > 0 C | 0.845±0.008 | 0.722±0.013 | 0.716±0.005 | 0.512±0.025 | 0.737±0.022 | 0.579±0.053 |
| > 5 C | 0.911±0.006 | 0.786±0.018 | 0.801±0.003 | 0.529±0.027 | 0.805±0.025 | 0.612±0.062 |
| > 10 C | 0.944±0.003 | 0.840±0.015 | 0.825±0.004 | 0.614±0.033 | 0.879±0.026 | 0.644±0.072 |
| > 20 C | 0.980±0.003 | 0.925±0.006 | 0.893±0.005 | 0.688±0.036 | 0.933±0.025 | 0.669±0.079 |
| > 30 C | 0.994±0.001 | 0.966± 0.005 | 0.934±0.006 | 0.731±0.039 | 0.961±0.027 | 0.681±0.087 |

**Table 3.** Regression model performance with Archaea, Bacteria, and combined dataset.

| Method | Archaea |  | Bacteria |  | Archaea + Bacteria |  |
| --- | --- | --- | --- | --- | --- | --- |
| | RMSE | $r^2$ | RMSE | $r^2$ | RMSE | $r^2$ |
| Random split | 8.06 ± 0.967 | 0.862 ± 0.024 | 6.76 ± 0.106 | 0.818 ± 0.005 | 7.31 ± 0.115 | 0.875 ± 0.003 |
| Phylogenetic distance split | 11.06 ± 1.005 | 0.772 ± 0.043 | 12.67 ± 0.862 | 0.370 ± 0.085 | 13.23 ± 1.643 | 0.590 ± 0.108 |

Results indicate that tRNA sequence alone can accurately predict both Archaea and Bacteria OGTs. Performance in all three datasets is highest for the randomly-split dataset, achieving 0.862, 0.818, and 0.875 *r*^2^ in Archaea, Bacteria and combined Archaea & Bacteria datasets, respectively (Table 3, Figure 3).

**Figure 3.**
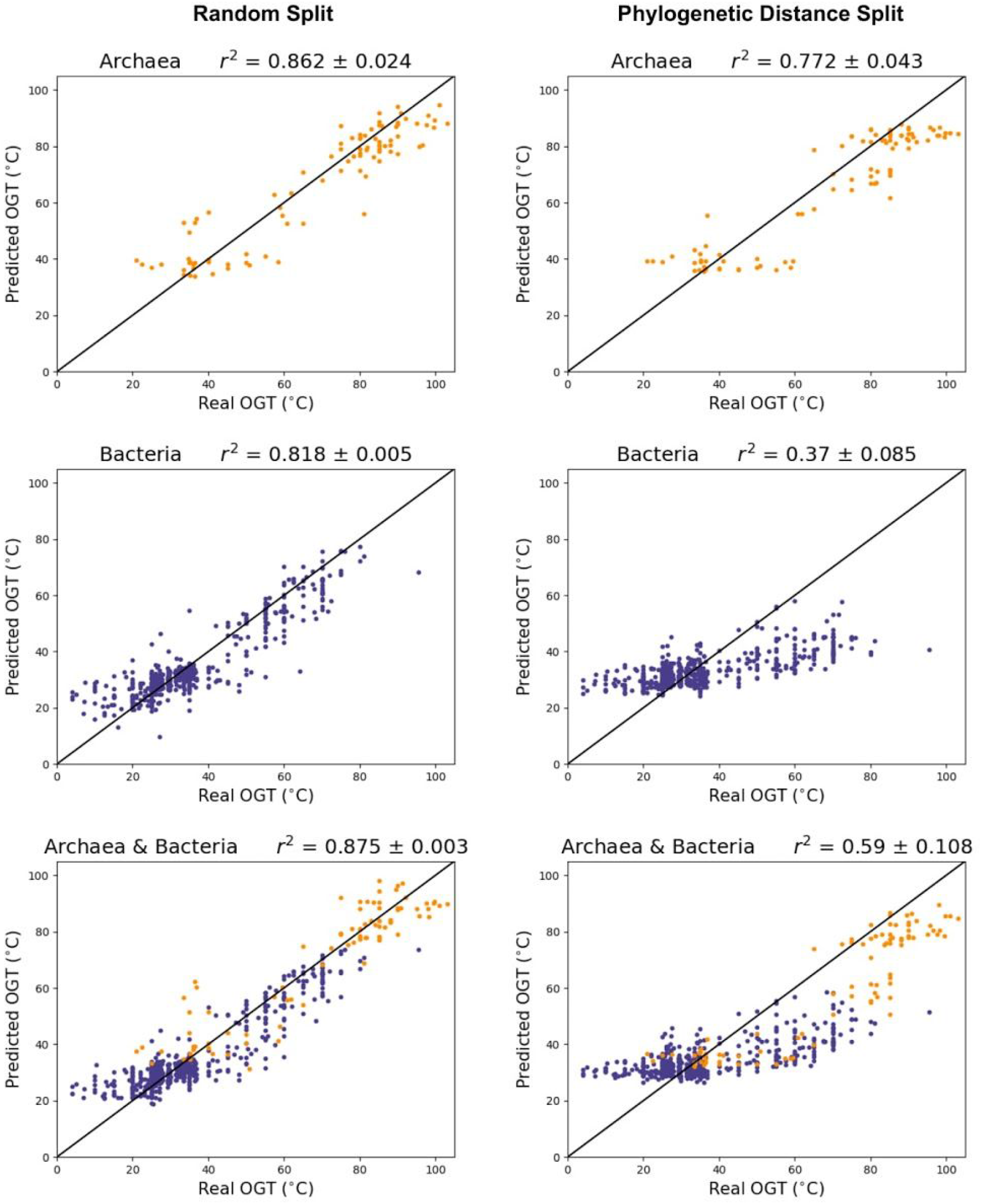
CNN model performance when data is split randomly (A-C) and split with phylogenetic distance (D-F). Purple = bacteria, orange = archaea.

One drawback of randomly splitting data into training and test sets is that similarity between individual observations in the training and test sets may lead to overly-optimistic model performances. When the end goal of a model is prediction, species relatedness is less of an issue. However, biological insights are more difficult to draw from a model that does not control for population structure within the dataset, since causal elements (in this case, causal nucleotides in the tRNA sequence) are confounded by structure due to shared evolutionary history. The phylogenetic distance model achieved 0.772, 0.370 and, 0.590 *r*^2^ in Archaea, Bacteria, and combined Archaea & Bacteria datasets, respectively (Table 3, Figure 3).

One criticism of convolutional neural networks is that they create a “black box” that can be difficult to interpret, making it hard to draw meaningful biological insight from a model. To try to understand what portions of the CNN matter for OGT prediction, we calculated attention statistics and evaluated the effects of directed mutagenesis on model predictions in bacteria. For every tRNA, the length of the stem and sum of CNN activation values was normalized to produce a single vector indicating the percent importance for the acceptor arm, D arm, anticodon arm, and T arm, respectively. The results show that the anticodon arm and T arm sequences are significantly more important for model predictions than the acceptor arm or D arm for both the random data split and for the phylogenetically-informed data split (Figure 3A, p < 2e-16).

### Regression model attention

To determine the importance of each stem structure for model predictions we selectively mutated 55 E. coli tRNAs to disrupt Watson/Crick base pairing. The results varied between the random-split and phylogenetic-split models. Mutations in the T arm of the tRNA led to a significantly different variance in OGT predictions for the random-split model, with some tRNA OGT predictions changing by 20 or 30°C (Bartlett test of homogeneity of variances, p=1.917e-15; figure 4), although the mean remained unchanged as did the mean and variance of predictions for other tRNA arms. In the phylogenetically-informed model, disrupting Watson-Crick base pairing did not affect model variances, but slightly increased the correlation between predicted minimum free energy of the tRNA structure and OGT predictions from −0.036 to −0.067 (p=3.5e-07).

**Figure 4.**
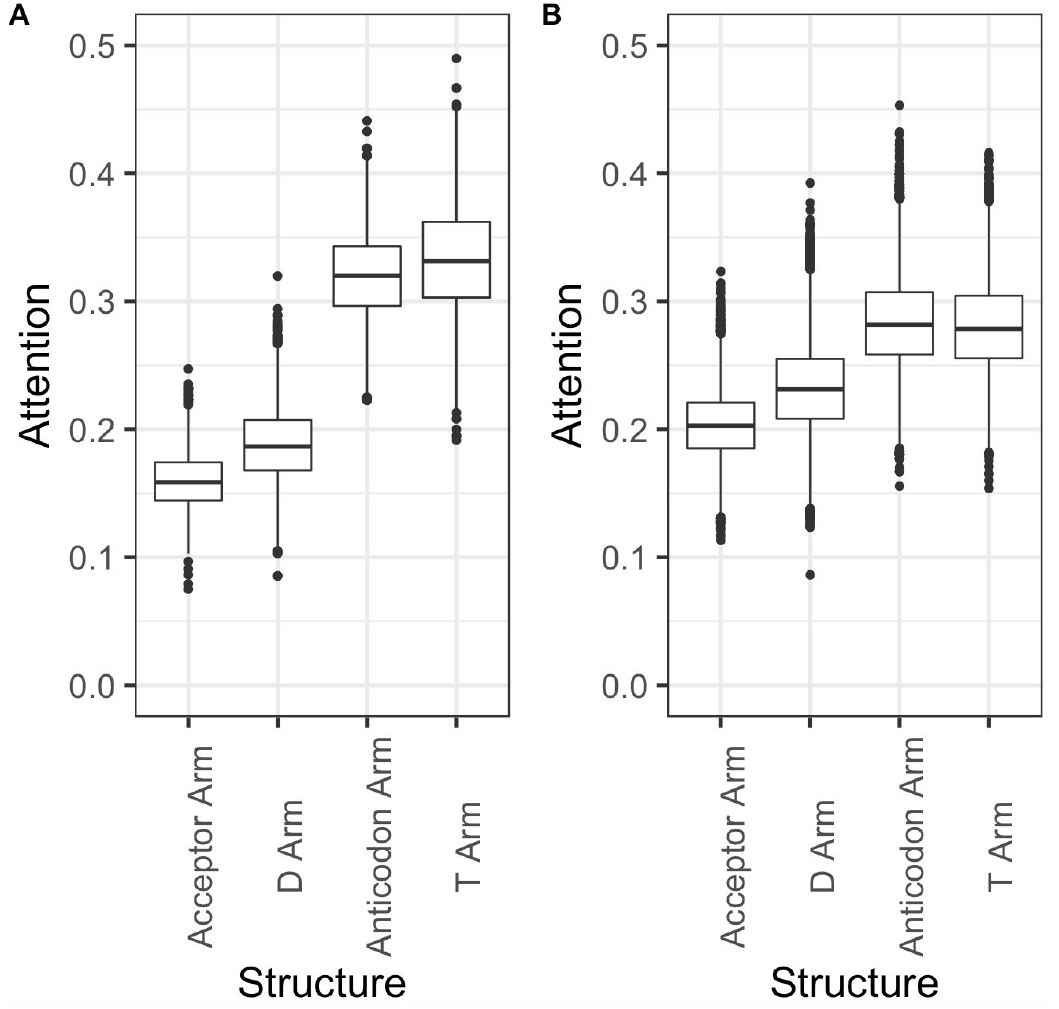
The percent importance assigned to each tRNA stem structure for OGT model predictions in A) Archaea and B) Bacteria. Both figures show model attention from the phylogenetic distance data split.

## DISCUSSION

### A machine learning model to predict prokaryote OGT

In this paper we discuss the application of machine learning models for predicting prokaryote optimal growth temperatures. Unlike previous models, we aimed to produce a model concentrating on only one element of cell biochemistry – the tRNA sequence – to predict OGT. An initial classification model was able to distinguish between pairs of tRNAs to identify which came from a species with higher OGT, suggesting that even individual tRNA sequences contain signatures of thermal adaptation. A second model uses the aggregate of effects from all tRNAs in an organism – together these sequences contain enough signal to predict organism OGT using a CNN regression model.

The classification model was able to classify species OGT from a single tRNA with an accuracy greater than 0.8 for all temperature differences above 10°C in models where data was split randomly. In all cases, models performed best when phylogenetic information was available to help with predictions, suggesting that species-specific differences in tRNA composition are indicative of tRNA thermal adaptation. This extra information about species relatedness was not available in the phylogenetic split model, resulting in lower classification accuracies. However, classification accuracies remained relatively high when classifying Archaea sequences, which may be due to the wider range of OGTs available for Archaea species.

Literature results using multiple linear regression to predict OGT achieved an *r*^2^ of 0.938 for archaea, 0.767 for bacteria, and 0.835 for a combined dataset by using genomic, tRNA, rRNA, open reading frames and proteome derived features and splitting training and test sets randomly (13). However, when only genomic and tRNA derived features are used by the authors, they achieved 0. 616 *r*^2^ (13). On the other hand, the proposed random-split CNN model achieves 0.875 *r*^2^ and shows a 42% improvement over literature *r*^2^ results for both the bacteria dataset and the combined bacteria and archaea dataset with only tRNA sequences as input. It is interesting that the CNN model outperforms in bacteria and in the combined dataset, but not in the archaea dataset. This may be due to the small number of archaea species with genome assemblies and OGT information available.

### Understanding model attention

The regression model attaches more importance to the regions of the input tRNA sequence that make up the anticodon and T arm structures, and mutating the structure of the T arm significantly changed the variance of model OGT predictions for the randomly-split dataset. Interestingly, both the T arm and anticodon arms are the regions of a tRNA at which post-transcriptional modifications are most concentrated. Although the regression model is not given information about post-transcriptional modifications, these modifications are often specific to a certain base and are known to affect tRNA folding and stability, especially in thermophiles (14). The fact that both the model with randomly-assigned training and test datasets and the model with phylogenetically-informed training and test datasets focus on the anticodon and T arms suggests that the CNN may be identifying signals related to post-transcriptional modifications in these regions. The T arm is also important for tRNA structure because its interactions with the D arm bend the tRNA into its appropriate three-dimensional structure. In its folded state, the T arm forms the elbow region of the three-dimensional tRNA structure. The elbow region is typically hydrophobic and is important for tRNA interactions with other RNAs and proteins (14).

Disrupting Watson/Crick base pairing in this region of the tRNA increases model OGT prediction variance, suggesting that the model may be recognizing the importance of this region for maintaining tRNA structure and function. Results differ slightly between the random-split model and the phylogenetic-split model. The increase in OGT prediction variance for mutations in the T arm suggests that the random-split model is looking at least partially for species-specific differences in this region. The increased correlation between minimum free energy of folding and OGT prediction suggests that the phylogenetic-split model is looking more at overall tRNA stability. However, the model is less accurate because it cannot use either information about species relationships or additional information about post-transcriptional modifications that would likely influence both MFE and species OGT.

The benefits of these models are threefold. First, the number of prokaryote sequences is growing, but additional information is often not available for these species, and developing culture protocols for new species can be challenging. Understanding likely OGT for new species is useful because it provides a starting point for labs wishing to develop culture protocols and further study these species. Knowing species OGT may also be useful in industrial processes requiring thermostable proteins, as this can provide insight into which species proteins are likely to be useful in such processes. Second, by using only the tRNA sequences we created a highly focused model that is independent of other cellular components. We use a minimum proportion of the overall genome sequence for predictions – only ~0.1% of total DNA in prokaryotes – to predict OGT. This is beneficial for downstream comparisons of temperature effects on protein, DNA, or other RNA features of the cell, as the OGT predictions from the tRNA model are independent of other cell components. Third, by using sequence data as direct inputs to the CNN model, we made use of automatic feature extraction and allowed the model to determine which tRNA features were most relevant. This removed researcher bias and did not require initial assumptions about which components of the tRNA sequence were most important.

### The importance of phylogenetic relationships between species

Although this model is able to accurately predict OGT within phylogenetic groups, it was unable to maintain high accuracy when predicting across phylogenetic groups, as demonstrated by the drop in prediction accuracy for the phylogenetic split in both the classification and regression models. These results indicate the importance of accounting for phylogeny when trying to extrapolate or draw biological insight from machine learning or other prediction models. In the current models, phylogeny is not part of the dataset, but the model clearly benefits from the shared evolutionary history between species in the dataset. Future studies may wish to determine how far insights from one group of organisms can be transferred – transfer may be limited to within species, clades, or superkingdoms, depending on the traits at hand. Additionally, other prediction methods, *i.e.,* multi instance learning algorithms, siamese neural networks can be applied to prediction problems in this study.

